# DelaySSAToolkit.jl: stochastic simulation of reaction systems with time delays in Julia

**DOI:** 10.1101/2022.01.21.477236

**Authors:** Xiaoming Fu, Xinyi Zhou, Dongyang Gu, Zhixing Cao, Ramon Grima

## Abstract

**Summary:** DelaySSAToolkit.jl is a Julia package for modelling reaction systems with non-Markovian dynamics, specifically those with time delays. These delays implicitly capture multiple intermediate reaction steps and hence serve as an effective model reduction technique for complex systems in biology, chemistry, ecology and genetics. The package implements a variety of exact formulations of the delay stochastic simulation algorithm.

**Contact:**, or

**Availability and Implementation:** The source code and documentation of DelaySSAToolkit.jl are available at https://github.com/palmtree2013/DelaySSAToolkit.jl.

## 1. Introduction

The stochastic simulation algorithm (SSA) is widely used to simulate the time-dependent trajectories for complex systems with Markovian dynamics (Gillespie, 1977). A major assumption behind these models is the memoryless hypothesis, i.e., the stochastic dynamics of the reactants is only influenced by the current state of the system, which implies that the waiting times between successive reaction events follow an exponential distribution.

Many common reaction systems encapsulate multiple intermediate reaction steps involving a large number of interacting species (Filatova et al., 2021). This leads to computationally expensive stochastic simulations using the SSA, which seriously limits the exploration of dynamics across large portions of parameter space. A viable alternative is the use of reduced models; while rigorous analytical reductions are possible if timescale separation exists (Mastny et al., 2007; Kan et al., 2016), more often than not it is more practical to replace multiple intermediate reactions by a single delayed reaction. This leads to a reduced model with non-Markovian dynamics, i.e. where the waiting times are non-exponential, which cannot be simulated by the conventional SSA.

Several exact simulation methods for reaction systems with delays were proposed (Barrio et al., 2006; Cai, 2007; Anderson, 2007; Ramaswamy and Sbalzarini, 2011). Some of these methods are available via software tools (Maarleveld et al., 2013; Barbuti et al., 2009). However, these packages offer limited functionality (implementing only few delay SSA algorithms and restricted to fixed delays, or mass action reactions) thus limiting the potential user base.

We present DelaySSAToolkit.jl, the first Julia package for modelling stochastic reaction systems with time delay. These delays can be either fixed or vary in time, i.e. each time a delay reaction fires, the delay is chosen from some user-defined probability distribution. It is also possible to model the case of heterogeneous time delays where there is a population of identical reactant systems, each of which has a reaction with a different but fixed time delay. As well, the toolkit can integrate simultaneous delay reactions and cascades of delay reactions using a variety of delay stochastic simulation algorithms. A comprehensive review of delay SSA theory and tutorials on using the software can be found at https://github.com/palmtree2013/DelaySSAToolkit.jl.

## 2. Materials and methods

In Figure 1, we summarize the workflow and showcase the software applied to an epidemic model, a birth-death stochastic model with heterogeneous delays (Cortez et al., 2021) and an auto-regulatory network of oscillatory gene expression with delays (Jiang et al., 2021).

**Figure 1:**
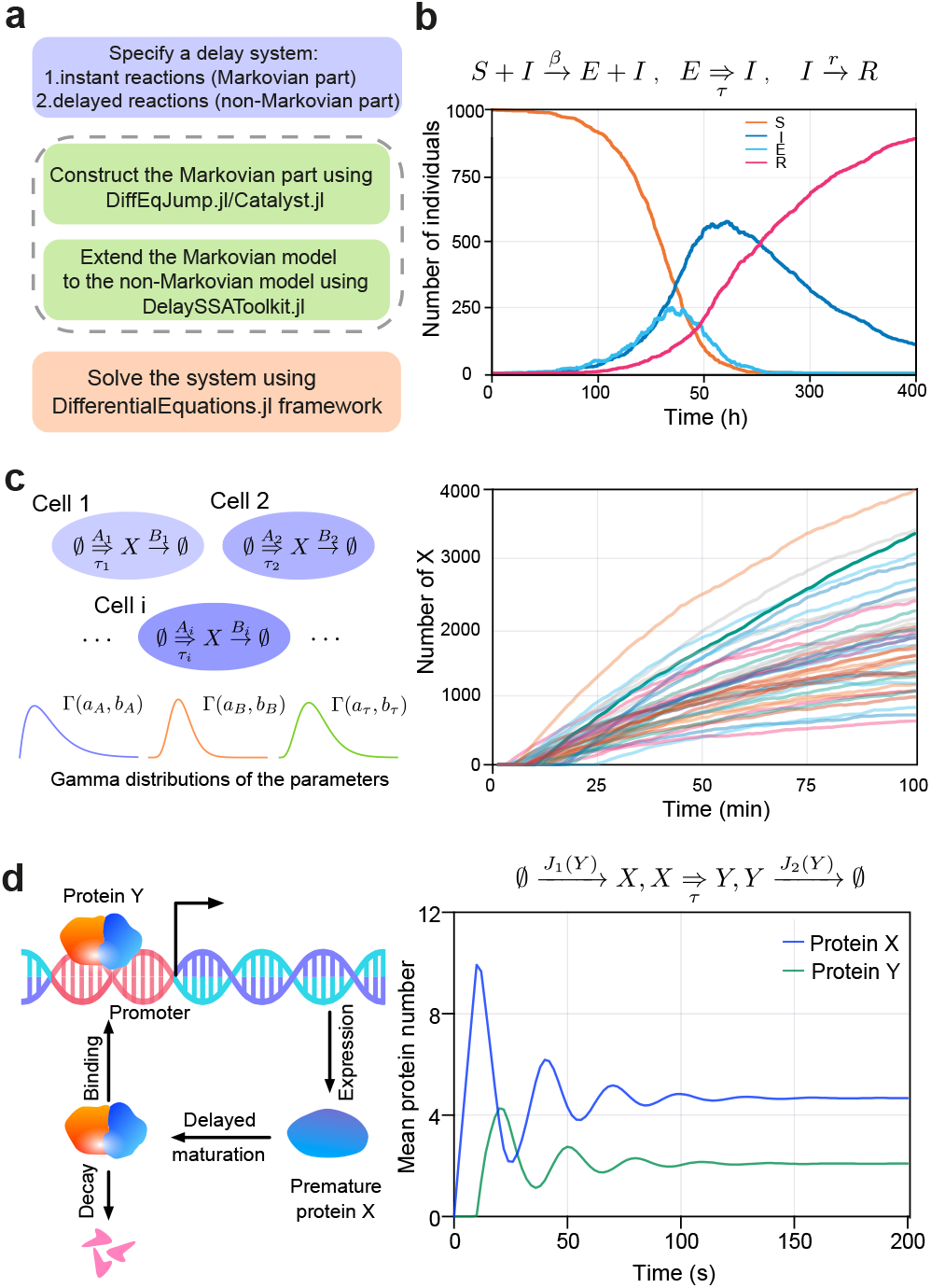
(a) The general workflow of stochastic simulation with delays in Julia using DelaySSAToolkit.jl and related packages. (b) An epidemic model with four compartments: susceptible (S), exposed (E), infected (I) and recovered (R): a susceptible contacted by an infected will firstly become an individual exposed to the disease and subsequently after a time delay *τ* becomes an infected individual which eventually recovers. The parameters are *β* = 10^−4^ h^−1^, *r* = 10^−2^ h^−1^, *τ* = 20 h. (c) Model of a population of cells with heterogeneous properties. For each individual, the production rate of a protein *X* is drawn from a Gamma distribution, i.e., *A* ~ Γ(8,0.23), the reaction is completed after a delayed time, *τ* ~ Γ(7,1). The protein degradation rate *B* follows Γ (9,625). We plot 40 simulated trajectories (each corresponding to a cell) subsampled every minute. (d) Model of an auto-regulatory negative feedback loop whereby a protein *X* is transcribed by a gene, transformed after a delay time *τ* into a mature protein *Y* which binds the promoter and represses the production of X. The propensities *J*_1_ (*ϒ*) and *J*_2_(*ϒ*) can be found in (Jiang *et al*., 2021). We also plot the oscillatory time-dependent mean values of two species with 10^4^ samples.

### 2.1. Model definition

The definition of a delay reaction system includes two parts, namely a Markovian part where the reactions affect the state of reactants instantaneously, and a non-Markovian part where the change of the state of reactants happens after a certain time delay *τ* (Figure 1a). The Markovian part of reaction networks can be constructed by specifying the reaction stoichiometry and the propensities using the Julia symbolic-numeric modelling framework embodied within Catalyst.jl (https://github.com/SciML/Catalyst.jl) and Model-ingToolkit.jl (Ma et al., 2021). Another way is via DiffE-qJump.jl (https://github.com/SciML/DiffEqJump.jl) by manually defining the stoichiometry and propen-sity of each reaction using a lower level programming interface.

DelaySSAToolkit.jl extends the Markovian models defined above to a delay system by specifying the causal relationships between the instant reactions and delayed reactions triggered by them. As such, DelaySSAToolkit.jl can handle a non-Markovian system containing any number of species and reactions with any type of smooth propensity functions with a wide range of delay types.

### 2.2. Delay SSA algorithms and solution handling

The algorithmic implementation of DelaySSAToolkit.jl is based on SSA provided by DiffEqJump.jl and the solution handling uses DifferentialEquations.jl. It currently supports four delay stochastic simulation algorithms: delay rejection method (Bratsun et al., 2005; Barrio et al., 2006), delay direct method (Cai, 2007), delay modified next reaction method (Anderson, 2007) and delay direct method with composition and rejection (Slepoy et al., 2008; Mauch and Stalzer, 2011). We note that the delay rejection and delay direct methods can offer better performance for small reaction networks, while the other two are preferable for systems with significant number of reactions. The computational efficiency of delay modified next reaction method and delay direct method with composition and rejection is improved for large-scale networks using the dependency graph and priority queue described in (Gibson and Bruck, 2000). Ensemble simulation is made easy using high-performance multi-threading/multiprocessing parallel computing interface implemented in DifferentialEquations.jl (Rackauckas and Nie, 2017), which also provides a number of numerical analysis and parameter estimation tools enabling even further study of the resulting delay system.

The resultant delay SSA solution can be given at user-specified time points or else at the exact event time points. The data structure of each solution is composed of three parts: the time points, the state of the reactants and the state of the delay channels at the corresponding time points. The recorded state of delay channels can be particularly useful in some cases, e.g. when modelling gene transcription, the delay reaction models the time between initiation and termination of transcription, and from the state of the delay channel one can reconstruct the positions of RNA polymerases on the gene (Fu et al., 2021).

## 3. Conclusion

DelaySSAToolkit.jl provides a feature-rich and userfriendly stochastic simulation tool for reaction systems with time delays. It is applicable to reaction networks of any size containing reactions with time/state-dependent propensity functions. Moreover, together with the other packages in the Julia ecosystem (Rackauckas and Nie, 2017; Ma et al., 2021; Sukys and Grima, 2021; Roesch et al., 2021), DelaySSAToolkit.jl makes the stochastic modelling of biochemical reaction kinetics efficient and accessible for Julia newcomers and experts alike.

## Acknowledgements

We thank Augustinas Sukys for useful discussions, feedback and independent testing of the Julia package.

## References

Anderson, D. F. (2007). A modified next reaction method for simulating chemical systems with time dependent propensities and delays. Journal of Chemical Physics, 127(21), 1716. _eprint: 0708.0370.

Barbuti, R., Caravagna, G., Maggiolo-Schettini, A., and Milazzo, P. (2009). On the interpretation of delays in delay stochastic simulation of biological systems. Electronic Proceedings in Theoretical Computer Science, EPTCS, 6(Comp-Mod), 17–29.

Barrio, M., Burrage, K., Leier, A., and Tian, H. (2006). Oscillatory regulation of hes1: Discrete stochastic delay modelling and simulation. PLoS Computational Biology, 2(9), 1017–1030.

Bratsun, D., Volfson, D., Tsimring, L. S., and Hasty, J. (2005). Delay-induced stochastic oscillations in gene regulation. Proceedings of the National Academy of Sciences of the United States of America, 102(41), 14593–14598.

Cai, X. (2007). Exact stochastic simulation of coupled chemical reactions with delays. Journal of Chemical Physics, 126(12), 297.

Cortez, M. J., Hong, H., Choi, B., Kim, J. K., and Josić, K. (2021). Hierarchical Bayesian models of transcriptional and translational regulation processes with delays. Bioinformatics, 38(1), 187–195.

Filatova, T., Popovic, N., and Grima, R. (2021). Statistics of nascent and mature rna fluctuations in a stochastic model of transcriptional initiation, elongation, pausing, and termination. Bull Math Biol, 83(1), 1–62.

Fu, X., Patel, H. P., Coppola, S., Xu, L., Cao, Z., Lenstra, T. L., and Grima, R. (2021). Accurate inference of stochastic gene expression from nascent transcript heterogeneity. bioRxiv.

Gibson, M. A. and Bruck, J. (2000). Efficient exact stochastic simulation of chemical systems with many species and many channels. Journal of Physical Chemistry A, 104(9), 1876–1889.

Gillespie, D. T. (1977). Exact stochastic simulation of coupled chemical reactions. The Journal of Physical Chemistry, 81(25), 2340–2361.

Jiang, Q., Fu, X., Yan, S., Li, R., Du, W., Cao, Z., Qian, F., and Grima, R. (2021). Neural network aided approximation and parameter inference of non-Markovian models of gene expression. Nature Communications, 12(1), 1–12.

Kan, X., Lee, C. H., and Othmer, H. G. (2016). A multi-timescale analysis of chemical reaction networks: Ii. stochastic systems. Journal of mathematical biology, 73(5), 1081–1129.

Ma, Y., Gowda, S., Anantharaman, R., Laughman, C., Shah, V., and Rackauckas, C. (2021). Modelingtoolkit: A composable graph transformation system for equation-based modeling. arXiv preprint arXiv:2103.05244.

Maarleveld, T. R., Olivier, B. G., and Bruggeman, F. J. (2013). StochPy: A comprehensive, user-friendly tool for simulating stochastic biological processes. PLoS ONE, 8(11).

Mastny, E. A., Haseltine, E. L., and Rawlings, J. B. (2007). Two classes of quasi-steady-state model reductions for stochastic kinetics. The Journal of chemical physics, 127(9), 094106.

Mauch, S. and Stalzer, M. (2011). Efficient formulations for exact stochastic simulation of chemical systems. IEEE/ACM Transactions on Computational Biology and Bioinformatics, 8(1), 27–35.

Rackauckas, C. and Nie, Q. (2017). DifferentialEquations.jl–a performant and feature-rich ecosystem for solving differential equations in Julia. Journal of Open Research Software, 5(1).

Ramaswamy, R. and Sbalzarini, I. F. (2011). A partial-propensity formulation of the stochastic simulation algorithm for chemical reaction networks with delays. Journal of Chemical Physics, 134(1).

Roesch, E., Greener, J. G., MacLean, A. L., Nassar, H., Rackauckas, C., Holy, T. E., and Stumpf, M. P. (2021). Julia for biologists. arXiv preprint arXiv:2109.09973.

Slepoy, A., Thompson, A. P., and Plimpton, S. J. (2008). A constant-time kinetic Monte Carlo algorithm for simulation of large biochemical reaction networks. Journal of Chemical Physics, 128(20).

Sukys, A. and Grima, R. (2021). MomentClosure.jl: automated moment closure approximations in Julia. Bioinformatics, 38(1), 289–290.

